# HGTector: An automated method facilitating genome-wide discovery of putative horizontal gene transfers

**DOI:** 10.1101/003731

**Authors:** Qiyun Zhu, Michael Kosoy, Katharina Dittmar

## Abstract

A new computational method of rapid, exhaustive and genome-wide detection of HGT was developed, featuring the systematic analysis of BLAST hit distribution patterns in the context of a priori defined hierarchical evolutionary categories. Genes that fall beyond a series of statistically determined thresholds are identified as not adhering to the typical vertical history of the organisms in question, but instead having a putative horizontal origin. Tests on simulated genomic data suggest that this approach effectively targets atypically distributed genes that are highly likely to be HGT-derived, and exhibits robust performance compared to conventional BLAST-based approaches. This method was further tested on real genomic datasets, including *Rickettsia* genomes, and was compared to previous studies. Results show consistency with currently employed categories of HGT prediction methods. In-depth analysis of both simulated and real genomic data suggests that the method is notably insensitive to stochastic events such as gene loss, rate variation and database error, which are common challenges to the current methodology. An automated pipeline was created to implement this approach and was made publicly available at: https://github.com/DittmarLab/HGTector. The program is versatile, easily deployed, has low requirements for computational resources, and is an effective tool for initial or standalone large-scale discovery of candidate HGT-derived genes.

## Introduction

Systematic studies have shown that horizontal gene transfer (HGT) is prevalent in prokaryotes (Abby et al., 2012; McDaniel et al., 2010; Ochman et al., 2000), where it serves as an important driving force of microbial evolution (Andam et al., 2010). HGT challenges the detection of vertical inheritance patterns in prokaryotes, and the application of conventional phylogenetic approaches to infer evolutionary history of microbial clades has seen increased limitations (Andam et al., 2010; Bapteste et al., 2009; Delsuc et al., 2005; Doolittle, 1999; Koonin and Wolf, 2009; Simonson et al., 2005). In essence, the ubiquitous nature of this process calls for the need to separate the vertical and horizontal patterns in evolutionary history of bacterial genomes. However, this is not straightforward in practice and is especially difficult for deep historical events because the horizontally acquired genes evolved along with the recipient genomes, gradually losing the signatures of their original hosts (amelioration) (Lawrence and Ochman, 1997). Furthermore, HGTs between closely related organisms, although common, are difficult to detect because in these cases donor and recipient share common compositional and phylogenetic features. So far, multiple computational methods have been developed to facilitate HGT detection, which may be loosely categorized into three main strategies based on sequence composition, phylogenetic analysis, or best BLAST matches (Doolittle et al., 2003; Zhaxybayeva, 2009; Zhaxybayeva and Doolittle, 2011). However, there appears to be poor agreement between outcomes of different methods, and comparative studies have repeatedly demonstrated that depending on method, very different sets of HGT-derived genes are identified from the same dataset, reflecting limitations in the current methodology for HGT prediction (Lawrence and Ochman, 2002; Ragan et al., 2006; Zhaxybayeva, 2009).

Given the rapid increase in available annotated genome data, and the associated computational challenge of analyzing such data, the BLAST best match method has remained a popular surrogate for first pass discovery analyses of gene histories that differ from the strict vertical pattern (Koonin et al., 2001). Specifically, this strategy is practiced by sorting BLAST hits by measures such as bit scores, an indicator of sequence similarity, and the best match organism represented by the top hit is identified for each gene (Altschul et al., 1997). If the best match is a distantly related organism, instead of one expected by vertical inheritance, then the gene is categorized as likely horizontally acquired (Eisen, 2000) (see Figure S1A, B). In practice, researchers often identify the best match using the criterion of bidirectional best hits (BBH) (Overbeek et al., 1999) to rule out potential paralogs. This approach has been applied by numerous studies (Charlebois and Doolittle, 2004; Clarke et al., 2002; Dagan et al., 2008; Koonin et al., 2001; Nelson et al., 1999; Smillie et al., 2011). Examples of programs featuring this approach include Pyphy (Sicheritz-Pontén and Andersson, 2001), PhyloGenie (Frickey and Lupas, 2004), NGIBWS (Charlebois et al., 2003), and DarkHorse (Podell and Gaasterland, 2007; Podell et al., 2008), although the latter also employs a user-definable filter threshold in combination with taxonomic scaling.

As expected, BLAST-based HGT detection has limitations. The bit score is based on sequence similarity, and provides only a rough estimate of the phylogenetic history between organisms. Also, best hits do not necessarily represent the nearest neighbors (Koski and Golding, 2001). Most importantly, odd patterns can also be caused by reasons other than HGT, such as lack of sequence information in related organisms (Lawrence and Ochman, 2002), gene loss events (Zhaxybayeva et al., 2007), stochastic similarity (Lawrence and Ochman, 2002) as well as database error (Willerslev et al., 2002). In practice, the predicted HGT candidates are often rejected by downstream phylogenetic analyses (Stanhope et al., 2001). Similar to phylogenetic HGT detection methods, BLAST best match is effective in detecting recent HGT events, but shows reduced sensitivity for ancient events, when donor and recipient sequences have already diverged over the long history of evolution (Boto, 2010). Additionally, merely a best match does not necessarily provide insights into the direction of gene flow.

Despite the listed issues, the best BLAST match methods make use of all sequence data available in GenBank. This sidesteps the need for manual subsampling and curation of comparative sequence data that is a challenging and time-consuming step in phylogenetic methods. This strategy is therefore likely to remain a feasible solution to explore microbial evolution in a first pass step, utilizing all of the ever increasing genomic data (Shokralla et al., 2012).

Considering this trend, we introduce a BLAST-based method to facilitate the detection of horizontal gene histories that aims to remedy some of the above outlined issues. This approach starts with standard all-against-all BLASTP, and is followed by an investigation of the weight distribution of all hits grouped by phylogenetically informed user defined categories (illustrated in Figures S1, 1 and 2). A general pattern of BLAST hit distributions (a fingerprint) of the genomes of interest is computed, and BLAST hit weights of each single gene are compared to the general fingerprint. This decreases sensitivity to stochastic disturbances. Because phylogenetic information is incorporated into this process, each resulting distribution is divided by uniquely defined cutoffs into typical and atypical gene populations. Using a combination of rules, a pool of genes that is putatively horizontally derived is reported. It is recommended that the atypical gene pool be subjected to downstream phylogenetic validation of HGT, which is implemented in this pipeline.

With this approach, the method retains all advantages of BLAST-based methods, such as rapid, efficient and exhaustive searches, which also facilitates re-analysis of data following the addition of new data. Rather than using general filter thresholds, and subsequent refining by taxonomic metrics (see LPI in DarkHorse (Podell and Gaasterland, 2007)), this method combines the two steps into one, and defines unique thresholds for each hierarchical level under consideration.

In order to assess the performance and robustness of this pipeline regarding the identification of putatively HGT-derived genes, it was applied towards simulated genomic data with known HGT events under consideration of various evolutionary forces. The method was also tested on real genomic datasets from multiple organismal groups, as exemplified by *Rickettsia*, whose HGT patterns have been previously studied (Blanc et al., 2007; Le et al., 2012; Merhej et al., 2011; Ogata et al., 2005; Weinert et al., 2009; Wolf et al., 1999). The consistency of these results was compared to those obtained by other methods, including sequence composition and phylogenetic approaches. Overlaps between results were investigated and discussed in the framework of technical and biological challenges behind each method.

## Methods

### Method overview

At the core of this approach is an all-against-all BLASTP search of the protein product of each protein-coding gene (referred to as gene hereafter) of the genome(s) of interest against the genome database. For each protein, the BLAST hits are recorded and sorted by their bit scores from high to low. The bit scores are then normalized by dividing them by the bit score of the query sequence against itself in order to account for the variety in lengths of proteins, so that every hit has a normalized bit score within the 0-1 range (Clarke et al., 2002). With the BLASTP search the organism corresponding to each hit and the taxonomic ranks of each organism are identified and recorded from the NCBI taxonomy database.

To proceed with the analyses, hits of each gene have to be divided into ***self***, ***close*** and ***distal*** groups. In other words, the pipeline doesn’t use phylogenetic trees or taxonomic lineage information directly, but rather allows the user to define three relational hierarchical categories, with the biological questions of the research in mind (Figures S1 and 1). This approach allows flexibility, because it can be scaled to the level of taxonomic/phylogenetic interest (e.g. intraspecies, intragroup, etc.), and it can be adjusted to frequent updates in bacterial taxonomy. The *self* group is considered the recipient, and always has to include the query genome(s), and, depending on analytical scale may also include its immediate sister organisms (e.g., different strains within the same species, or different species within the same genus). The *close* group will include representatives of the putative vertical inheritance history of the group (e.g., other species of the same genus, or other genera of the same family which the query genome belongs to). The *distal* group includes all other organisms, which are considered phylogenetically distant from the query genome (e.g., other families, orders, etc.). The method will then aim to identify genes that are likely derived from directional gene flow from groups of organisms within the *distal* group to members of the *self* group.

We introduced a measure to quantify quality of BLAST hits of each group. This measure (“weight”) is calculated per group by summing up the normalized BLAST bit scores of hits. Because three categories were defined a priori, this step will result in three weights per gene. The three weights of all genes of the query genome are considered as three independent statistical populations. If multiple genomes are analyzed together, the weight populations can be merged. The three distributions together are defined as a fingerprint of the input genome(s).

A cutoff is set up to divide the weight distribution of each group (*self*, *close* and *distal*) into typical (larger than or equal to cutoff value) and atypical (smaller than cutoff value). If most genes of the genome(s) have a vertical inheritance history, the typical portion should include a majority of the genes, while the atypical portion should only include a small, but significant subset of genes whose hits of this specific group are underrepresented. The cutoff defines the stringency of prediction: the higher the weight cutoff, the more genes are considered as being atypical. The pipeline implements statistical approaches to compute the cutoffs, but the user is free to implement and use their own statistics. Because each genome or set of genomes generates a different fingerprint, cutoffs will vary, and are not transferrable across analyses.

For any gene to be predicted as putatively horizontally acquired, the following rules apply, which take into account all three weight distributions for that gene:

**Rule 1:** The gene is below cutoff (atypical) in the *close* weight distribution, suggesting that the orthologs of the query gene are absent in all or most of the sister groups of the organism of interest. This means that the BLAST hits belonging to this group are significantly underrepresented, in terms of number of hits or bit score, or both (Figure S1B-D). This phenomenon can be explained by: (i) the gene lacks a vertical inheritance history, or alternatively, (ii) the gene was vertically inherited, but underwent multiple independent gene loss events in the sister groups, a case that is usually less likely to be true (Doolittle et al., 2003). In the ideal scenario of a high likelihood of HGT, the weight should be zero, meaning that there is no close hit (Figure S1B, C). In real datasets however, there are sometimes sporadic close hits with low bit scores (Figure S1D). These hits may be due to stochastic sequence similarity, secondary HGTs, paralogous genes or other mechanisms. This situation typically causes false negatives in conventional best BLAST match methods (Doolittle et al., 2003). However, in the present method, these sporadic hits will not significantly alter the overall weight of a gene, thus hardly affecting the prediction results.

On the other hand, for a vertically transmitted gene, its orthologs may not always be present in each and every sister lineage. Occasional gene loss events may take away some of the expected number of *close* hits (Figure S1E). This situation is a major source of false positives in best BLAST match methods (Zhaxybayeva et al., 2007). However, this problem is effectively overcome in the present method, because sporadic absences of hits do not make the overall weight atypical. Moreover, one or a few high-score *distal* hits caused by natural (outgoing HGT from the *self* group to the *distal* group) or artificial reasons (contamination, mislabeling, etc.) (Figure S1F) can easily deceive conventional methods (Willerslev et al., 2002). However, the present method is immune to this problem, as the *close* weight remains unchanged in this situation.

**Rule 2:** The gene is equal to or above cutoff (typical) in the *distal* weight distribution, meaning that the hits from distant organisms are not underrepresented (Figure S1A-F, H). This criterion sets up a filter at the donor side of potential HGT events: given the gene was transferred from a representative or ancestor of organisms that belong to the *distal* group, BLAST hits of the *distal* group itself should not be underrepresented. Otherwise, it is not convincing to conclude that the gene is HGT-derived.

The goal of this rule is stringency, in order to better distinguish putative HGT events from other scenarios that can also make a gene’s *close* weight atypical: (1) de novo gene origination within the *self* group, (2) inaccurate genome annotation that considers a noncoding region a gene, or (3) HGT from an unsequenced organism that is not phylogenetically close to any sequenced groups of organism. These scenarios usually result in few to no *distal* hits (Figure S1G, upper panel). Meanwhile, (4) sequence similarity due to randomness instead of homology may also bring in some *distal* hits with low bit scores (Figure S1G, lower panel).

**Rule 3 (optional):** The gene is below cutoff (atypical) in the *self* weight distribution. It means that a gene is only sporadically detected, rather than being prevalent across the whole *self* group of organisms (Figure S1B, H). This option will restrict the prediction to a subset of putatively HGT-derived genes that were acquired by specific lineages of the *self* group, instead of the whole *self* group. Often, these could be recent transfer events.

To assess the source of a putative HGT event, the best match organism from the *distal* group is reported. It is important to understand that this best match is likely not the actual physical donor, but may be an extant representative of an ancestral, extinct donor. We recommend using “donor link” to describe the directionality of transfer, and relationship between these organisms, instead of “donor” (Merhej et al., 2011).

A flowchart of the procedures of this method is illustrated in Figure 1.

**Figure 1.**
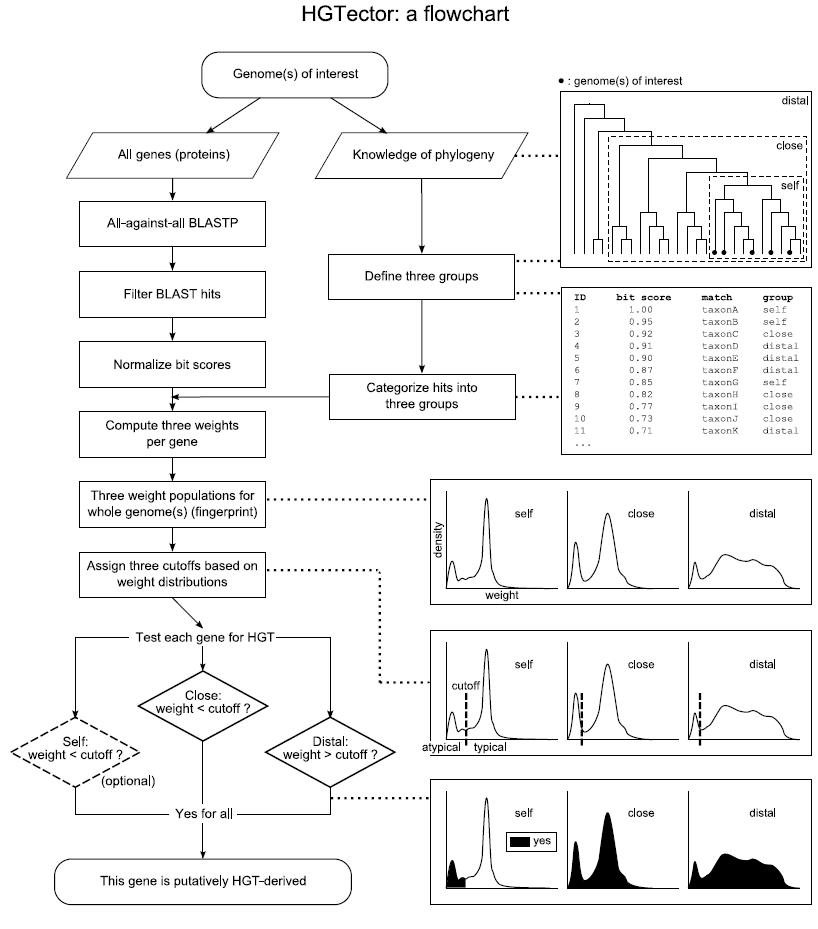
Flowchart illustrating the procedures of the HGTector method. Parallelograms indicate input data or information, rectangles indicate processing steps, diamonds indicate decisions, and rounded rectangles indicate start and end of the work flow. Graphic illustrations of hypothetical phylogenetic tree, BLAST hit table and fingerprint are drawn on the right side of their corresponding steps.

### Computational pipeline

#### General procedure

The HGTector pipeline (publically available at: https://github.com/DittmarLab/HGTector) is written in Perl, and is cross-platform supported, running in Windows, Mac OS and Linux systems. It requires only the Perl interpreter with its core modules, a default component of most Mac OS and Linux distributions and is very easy to install in Windows systems. In the default mode, the program depends on no additional software or local databases to run. This characteristic maximizes the ease of installation for users without professional computer background and resources. Optionally, it calls R (R Core Team, 2013) to perform advanced statistical computing and graphing. Parameters of the program are managed by a central configuration file, which can be created and edited manually or via a graphical user interface (GUI). The program is composed of the following procedures:

First, the script performs batch BLASTP of protein products of multiple genes supplied by the user. Multiple formats ranging from simple lists of NCBI accession numbers to annotated genomes in GenBank format are supported. It runs BLASTP either via web connection to the NCBI server, or with a standalone BLAST program and a local database. It also harvests taxonomic information of each hit for each gene from the NCBI taxonomy database. Hierarchical taxonomic reports (NCBI TaxBlast) and sequences of hits (original or aligned) can be retrieved optionally. BLAST results are saved in NEXUS format (Maddison et al., 1997), which can be directly viewed by text editors, or opened as multiple sequence alignments by external programs such as SeaView (Gouy et al., 2010), for additional analyses. This characteristic facilitates downstream analyses, and compatibility with other programs.

BLAST hits are filtered by multiple optional functions to overcome putative taxon sampling bias that may affect BLAST hit distribution: (1) When multiple hits are present for one organism (e.g., dozens of copies of a mobile element), only the best hit is maintained, representing the putative ortholog of the query protein (Tatusov et al., 1997). (2) When multiple genomes are present for one species (e.g., hundreds of sequenced *Escherichia coli* strains), only the genome that contains the best hit is maintained. (3) Taxonomic name keywords or IDs that represent unwanted BLAST targets, such as inaccurately defined taxonomic ranks (e.g., genus *Clostridium*) and biological categories (e.g., “environmental samples”), can be specified so that these hits will be omitted.

By default, the program will exclude genes without any non-*self* hits from subsequent analyses, because they may represent ORFans (Fischer and Eisenberg, 1999), resulting from de novo gene origination events (which are very rare (Long et al., 2003)), or transfer events from unknown sources that are very dissimilar from any sequenced genomes. Alternatively, they may represent genome annotation errors, which have been long recognized as a common and perturbing issue (Brenner, 1999; Kohane et al., 2012; Wong et al., 2010). While these genes are not considered in the subsequent analysis, the genes are reported as “putative ORFans or annotation errors”, or POE, in this pipeline (Figure S1G, upper panel). This allows the user to check which POEs were omitted, and if necessary make further adjustments to the analytical set up.

Based on the retrieved taxonomic information, the program can automatically formulate a grouping scenario, in which the lowest rank that includes all input genomes is defined as the *self* group, and the next higher rank as the *close* group. Because taxonomic classifications in GenBank may not always reflect the most current natural groupings of organisms, users may manually define hierarchies based on current knowledge of phylogeny and the purpose of their research.

With a properly defined grouping scenario, the program then calculates the three weights of each gene, computes a fingerprint of the whole genome(s), defines proper cutoffs and determines the population of atypical events, and possible HGTs based on the selected rules. Basic statistical parameters of the three distributions of weights, as well as the weight populations themselves are reported. The fingerprint may be visualized by box plots, histograms, density plots and scatter plots (Figures 2 and 4). Statistical analyses and graphing of BLAST hit distributions are automated in the program. They are performed using Perl codes, or by sending commands to R (R Core Team, 2013). The communication between Perl and R is utilized by the Perl module “Statistics::R” . While multiple statistical approaches are available for the user’s choice, the typical procedures, which are used in the tests described in this article, are as follows:

**Figure 2.**
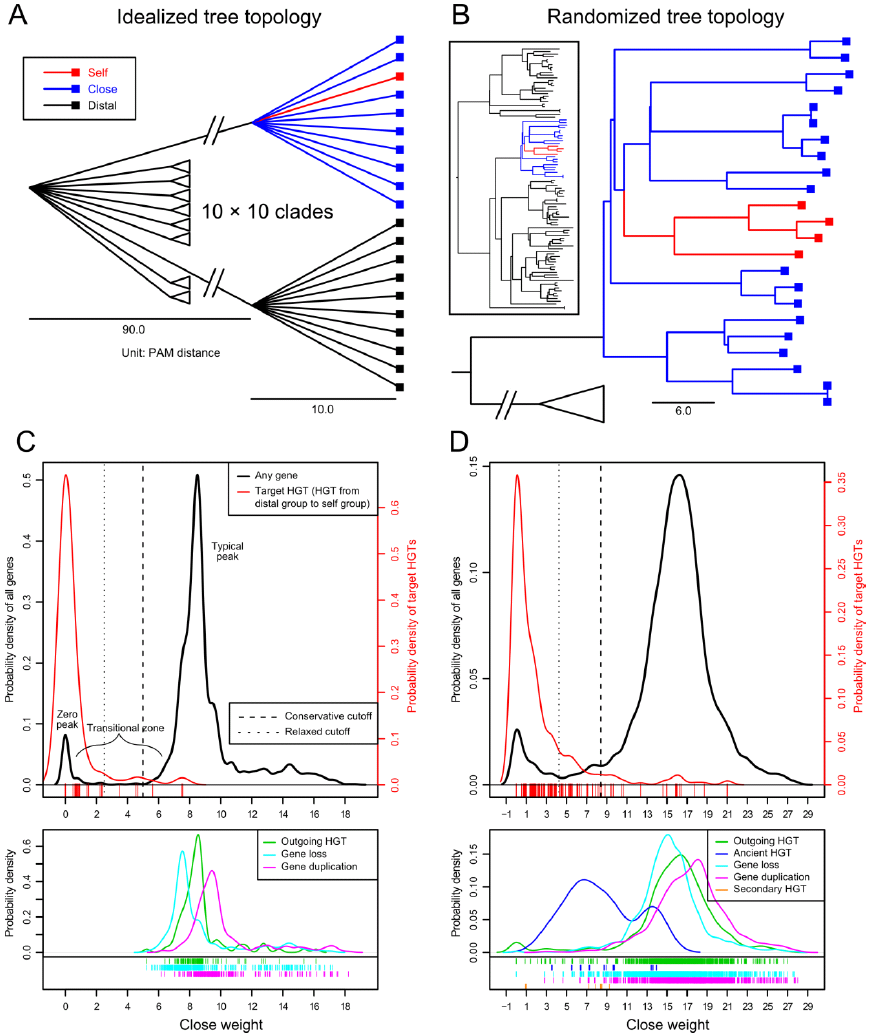
Tree topology and fingerprint (distributions of BLAST hit weights) of tests on simulated genomes. One representative test using either the idealized tree topology (**A**) or the randomized tree topology (**B**) is depicted (see text). Kernel density function of *close* weight distribution for both topologies (**C, D**), shows the distribution of all genes in the input genomes in black, and that of actual positive genes (derived from HGT events from *distal* group to *self* group) in red. Genes involved in other simulated evolutionary events are shown in different colors in the lower panels. Locations of these genes in the general distribution are indicated as a rug below each plot. The scales of x-axes between the upper and the lower panels are identical. The cutoffs computed by the program that distinguish the atypical region from the typical region are represented in dashed (for relaxed criterion) and dotted (for conservative criterion) lines.

#### Cutoff determination

The program performs kernel density estimation (Silverman, 1981) to obtain a function of probability density distribution of the *close* weight for all genes. By default it uses Gaussian kernel smoothing with Silverman’s rule-of-thumb bandwidth selector (Silverman, 1986). The user is allowed to choose a proper bandwidth selection factor that controls the smoothness of the curve. The function is plotted and made visible to the user in real time. Statistically significant local minima (pits) and maxima (peaks) are computed using the “pastecs” package (Frederic et al., 2013) with default parameters following Kendall’s information theory (Kendall, 1976), and their x-coordinates are recorded and displayed to the user. The program then automatically identifies a local minimum separating the typical from the atypical proportion of the gene population under consideration. Specifically, the biggest peak or group of continuous peaks in terms of number of genes it covers is identified as the typical region, and the rest is defined as the atypical region (Figures 2 and 4).

In addition to the apparent typical peak, there is usually a clearly identifiable peak located close to zero (referred to as the “zero peak” hereafter). This peak usually includes “ideal” putatively HGT-derived genes (= BLAST hit of zero). Between the zero peak and the typical peak is a transitional zone that likely consists of genes with ambiguous evolutionary history (see Results and Discussion).

The program automatically reports two cutoffs: the x-coordinate of the identified local minimum is naturally chosen as a cutoff (referred to as the “relaxed cutoff” hereafter). However, based on results of repeated tests, we recommend using the second cutoff, which is defined as the arithmetic mean of the x-coordinates of the local maximum of the zero peak and the local minimum of the selected pit (referred to as the “conservative cutoff” hereafter) (Figure 2C, D). The choice between the two cutoffs depends on the goal of research, but for the identification of putative HGT events, the conservative cutoff is preferred as it meets a balance between precision and recall (see Results and Discussion).

The program also implements several functions to assess the statistical significance of separating atypical genes from typical ones. For the whole weight population, the program performs Hartigans’ dip test (Hartigan and Hartigan, 1985) to assess the non-unimodality of the weight distribution, which essentially is the statistical significance that a distribution can be divided into two or more distinct parts. The test is performed by calling the “diptest” package in R (Maechler and Ringach, 2009). The dip statistic and the p-value for the test for unimodality are reported. For each individual gene, the program computes a density-based silhouette (dbs) (using the “pdfCluster” package in R), which is a statistical measure of confidence that a certain data point (gene) is allocated to a cluster (here the atypical region) (Menardi, 2010).

The same procedures apply to the distribution of the *distal* weight (and the *self* weight, if the optional rule 3 is chosen). In addition to the above described statistical approaches, power users may also perform extra statistical analyses based on the weight data output by the program, and type user-defined cutoffs.

Based on the cutoffs, the program reports a population of genes with an atypical, non-vertical history, which in the context of the a priori provided phylogenetic information represents a putative horizontal history. The results are summarized in a choice of plain text, web page (HTML) or Excel spreadsheet formats. The latter allows for convenient downstream statistics of outputs, and includes hyperlinks that allow users to track each of the genes back to their original BLAST report. It not only reports the number and percentage of putatively HGT-derived genes, but also optionally categorizes each gene in three contexts: (i) By putative donor group, which is described by user-designated higher taxonomic rank of the best match organism (based on GenBank annotations). (ii) By functional annotation of protein products, which is provided by external sources, such as the output of Blast2GO (Conesa et al., 2005). (iii) By gene orthology (evolutionary history of each individual gene family across input genomes), which is identified by a built-in function of BLAST hits clustering or from external sources, such as the output of OrthoMCL (Chen et al., 2006). These reports allow users an intuitive view of the prevalence of HGT-derived genes and the evolutionary, ecological and functional implications of HGTs at levels of individual genes, gene families, whole genomes and multiple phylogenetically related genomes.

The whole analysis can be performed on a regular personal computer, as it does not require extensive computing power. Batch-BLAST is the most time-consuming step, which typically lasts several hours to several days, depending on the number of protein-coding genes in the genome(s) of interest. The statistical analysis typically takes only minutes. Data generated by the program can be parsed and reused by other programs for multiple purposes.

As an additional, and important function, the program provides a complete phylogenetic pipeline, which automates the process of multiple sequence alignment, alignment trimming and phylogenetic tree reconstruction, by calling external local programs such as ClustalW (Thompson et al., 1994), Gblocks (Castresana, 2000) and RAxML (Stamatakis, 2006), and parsing their outputs. Reconstructed phylogenetic trees are annotated with organismal names and are attached to BLAST reports, which in turn can be directly viewed by external programs such as FigTree (Rambaut, 2013). This function allows users to monitor and validate prediction results by manually checking the evolutionary scenarios of individual genes.

### Analysis of simulated genomic data

To assess the performance of this method under the impact of various evolutionary scenarios, as well as to compare HGTector to conventional BLAST methods, we tested the above-described pipeline on simulated genomic data.

#### Simulation of genome evolution

Simulated genomic data were generated by ALF (Artificial Life Framework) version 1.0, a program that simulates genome evolution (Darriba et al., 2012). In each simulation, 100 species evolved from one randomly generated root genome containing 1000 protein-coding genes that are no shorter than 50 aa. During the process random inter-genomic HGT events occurred under a pre-defined global rate, which varied between simulations (see below). The process also simulated the following evolutionary forces in addition to HGT, at random rates: speciation, character substitution, insertion and deletion, GC-content amelioration, rate variation among sites and among genes, gene duplication and loss. Many of these forces are known to affect HGT prediction (Lawrence and Ochman, 2002; Willerslev et al., 2002; Zhaxybayeva et al., 2007).

Two types of simulations were performed. First, an idealized, pre-defined tree topology was used, in which all representatives of *close*- and *self*-designation are grouped in a polytomy, signifying (in this case) equal genetic distance, thus eliminating the impact from evolutionary artifacts. The rest of the taxa are placed relatively distant (unrelated) in the tree to lower the possibility of stochastic BLAST matches. The topology is detailed as follows: 10 clades branch from the tree base simultaneously at time point 0 (unit: PAM distance, same below). In each clade, 10 species branch off simultaneously at time point 90. The tips are at time point 100 for all species. For each clade, one species was randomly chosen as the input genome (also the *self* group), while nine species were defined as the *close* group. All 90 species in other clades are considered as the *distal* group (Figure 2A). All ten clades were analyzed in this manner. This simulation was replicated 10 times (= 100 analyses), with the HGT rate ranging from 0 (negative control) to 0.0045 with an interval of 0.0005.

Second, a randomized birth-death tree was generated in ALF per simulation (birth rate = 0.1, death rate = 0.01, height = 1000), mimicking a more realistic topological situation. A random clade was manually chosen from the tree, as long as it met the following criteria: (1) 3-8 *self* species that formed a clear monophyletic group; (2) 10-20 *close* species that were closely related to the *self* clade; (3) the *self* and *close* species together formed a clear monophyletic group that was independent from all other species (the *distal* group) (Figure 2B). This simulation was replicated for 100 times, with the HGT rate of each replicate randomly sampled from a range of 0 to 0.005.

Evolutionary events (HGT, gene duplication, gene loss, etc.) simulated in each analysis were extracted from the ALF log file. The time, species and genes involved in these events were recorded. HGT events involving the *self* group were further categorized by their donor, recipient and time into the following groups: target HGT (HGT from the *distal* group to the *self* group, which are the actual positives to be targeted by HGTector), ancient HGT (HGT from the *distal* group to the ancestor of the *self* group), outgoing HGT (HGT from the *self* group to the *distal* group, which is equivalent in consequence to a database error that mislabels a sequence with another organism), and secondary HGT (target HGT combined with one or more additional transfers between the *self* and the *close* groups). Gene loss and gene duplication events taking place in the *close* group were singled out as they directly influence the distribution of the *close* weight.

For each simulation (idealized or randomized tree topologies), a BLASTP database including the protein sequences from all 100 genomes was created using the standalone BLAST program (Altschul et al., 1997). All-against-all BLASTP was performed with an E-value cutoff at 1×10^−5^ for all genes in the selected *self* genomes, considering at least 20 hits. In order to demonstrate the effect of gene duplication on the prediction result, the program option of excluding paralogs was turned off.

A modified version of the HGTector pipeline was created to parse the ALF outputs and to perform analysis. Both the conservative and the relaxed cutoff criteria were tested. The fingerprint was plotted along with the distribution of actual positives (target HGTs) as well as other evolutionary scenarios (see above) for manual observation in addition to statistical analysis.

#### Comparison to conventional BLAST approach

The performance of HGTector in the context of identifying atypical and putatively HGT-derived genes, was compared against that of commonly used conventional BLAST-based methods by modifying the pipeline to mimic the conventional approach, which does not consider overall hit distribution. Specifically, we considered HGT-events for the scenario where no *close* hits are present (criterion: C=0); or the scenario where there is at least one *distal* hit with a bit score higher than those of any *close* hits (criterion: D>C).

#### Assessing Performance

The predicting power in the context of target HGTs under each criterion, as well as for conventional BLAST was assessed by precision and recall. Precision describes the number of true positives over the number of all predicted positives (= How many of the predicted HGTs are real), and recall describes the number of true positives over number of all actual positives (= How many of all real HGTs were found), which are commonly used performance markers for binary classifications. Statistical analysis and plotting were conducted using R (R Core Team, 2013).

### Application of HGTector to real datasets

HGTector was further tested on a variety of real-world genomic data, covering bacteria (5), unicellular eukaryotes (1) and human (1) (Table 1), the last of which serves as a realistic negative control since HGTs into genomes of higher eukaryotes are known to be very rare (Doolittle, 1999). The most in depth analysis was conducted on the *Rickettsia* dataset, because it affords comparisons to previous results obtained with a variety of methods (Langille et al., 2008; Le et al., 2012; Merhej et al., 2011; Ogata et al., 2005; Podell et al., 2008).

**Table 1.**
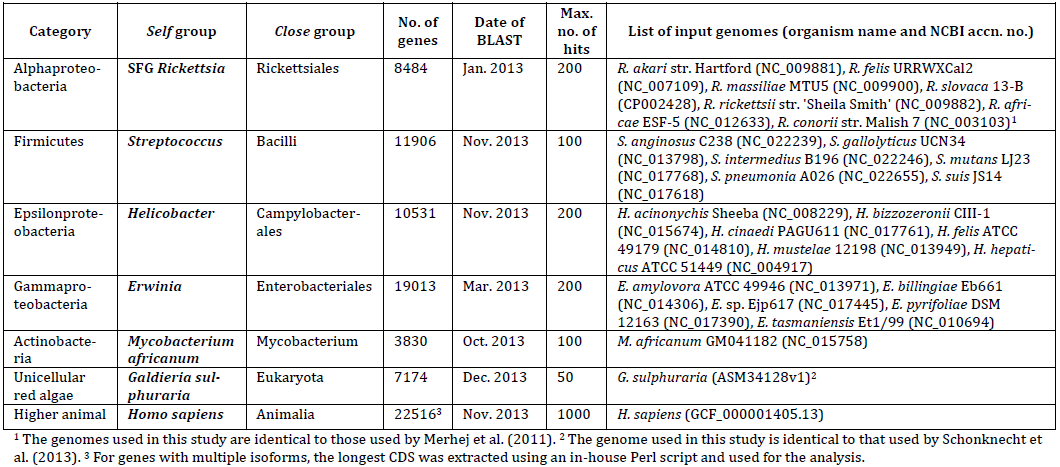
Real genomic datasets tested in this study

#### Analysis on the Rickettsia dataset

Out of all available *Rickettsia* genomes, we selected seven representative *Rickettsia* species with fully-annotated genomes for this analysis (Table 1). All of these species belong to the spotted fever group (SFG), a traditional classification group of *Rickettsia*. A grouping scenario was chosen based on the taxonomy and phylogeny of major *Rickettsia* species, which has been well resolved by recent studies (Merhej et al., 2011; Vitorino et al., 2007; Weinert et al., 2009). Specifically, we defined the *self* group as SFG (NCBI taxonomy ID: 114277), the *close* group as order Rickettsiales (766), excluding SFG, and the *distal* group as all non-Rickettsiales organisms.

All-against-all BLASTP was performed against the NCBI non-redundant protein sequences (nr) database with an E-value cutoff at 1×10^-5^. A soft filtering for low sequence complexity regions, which was suggested as the optimal parameter for ortholog identification (Moreno-Hagelsieb and Latimer, 2008), was used. Hits with organism names containing these words were excluded for the purpose of this analysis: unknown, uncultured, unidentified, unclassified, environmental, plasmid, vector, synthetic, and phage. Up to one hit from each organism was preserved. A maximum number of 200 hits were preserved for each protein. A global fingerprint was computed and graphed to describe the pattern of BLAST hit distribution of all seven genomes. Cutoffs for the three groups of weights were computed using the built-in kernel density estimation function under the conservative criterion. The default rules 1 and 2 were applied.

#### Assessing stochastic events using real datasets

To test the impact of stochastic events on prediction results, the following simulations were carried out on the *Rickettsia* dataset: (1) database error (some sequences are mislabeled by incorrect organism names), (2) gene loss in the *close* group, (3) rate variation in the *close* group, (4) rate variation in the input genomes, and (5) taxon sampling bias (some groups of organisms are overrepresented in the genome database). Multiple degrees of modification intensity (*x*) for each type of events were tested, each having 100 replicates (except for taxon sampling bias). Specifically:

1. Database error. For each hit, there is *x* proportion of chance that its corresponding organism was assigned to an organism randomly sampled from the pool of BLAST results of all genes.
2. Gene loss in the *close* group. For each *close* hit, there is *x* proportion of chance of being removed from the BLAST hit table.
3. Rate variation in the *close* group. For each *close* hit, there is *x* proportion of chance that its bit score is divided by a factor subject to a gamma distribution with shape parameter *k* = 2 and scale parameter *θ* = 1:

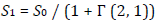

Where *S*_0_ and *S*_1_ refer to the bit score before and after manipulation, respectively.
4. Rate variation in the input genomes. For each query gene, there is × proportion of chance that the bit scores of all its hits are divided by a factor subject to a gamma distribution same as above.
5. Taxon sampling bias. For selected representative groups of organisms from *close* and *distal* groups (see Results and Discussion), all hits belonging to this group were replicated into *x* copies (*x* identical but separate hits).

HGTector analysis was conducted on these replicates using the same procedures as in the standard analysis on the unmodified dataset (see above). The results were compared to the results derived from a conventional BLAST-based approach under the D>C criterion (see Analysis of simulated genomic data). Precision and recall of the results were computed using the result of the standard analysis as the reference.

#### Cross-method comparison

In order to evaluate the performance of HGTector on the Rickettsia dataset in the context of other available methods, results were compared to two examples from each of the three currently employed strategies: BBH (bidirectional best hit) (Overbeek et al., 1999) and DarkHorse (Podell and Gaasterland, 2007; Podell et al., 2008) based on best BLAST match; GIST (Hasan et al., 2012) and IslandViewer (Langille and Brinkman, 2009) based on sequence composition, and two studies conducted by Merhej et al. (2011) and Le et al. (2012), using phylogenetic approaches. We exemplified this comparison on the *R. felis* genome, which previously has been demonstrated to have high HGT frequency (Merhej et al., 2011; Merhej and Raoult, 2011; Ogata et al., 2005).

Specifically, BBH analysis was performed as a built-in function of the present pipeline. This method is similar to the D>C criterion as described above, except for an additional reverse BLAST step with the same parameters to confirm that the two sequences are each other’s best match within their host genomes. The result by DarkHorse was downloaded from the DarkHorse server (darkhorse.ucsd.edu). Default parameters were used, in which the BLASTP E-value cutoff is also 1×10^−5^. All three available phylogenetic granularities, strain, species and genus, were used and the results were merged, in order to maximize the discovery rate. Both GIST and IslandViewer are targeting large pieces of heterogeneous genomic regions, or genomic islands (GI) (Hacker and Kaper, 2000). GIST is a synchronization of five subprograms: AlienHunter (Vernikos and Parkhill, 2006), IslandPath (Hsiao et al., 2003), SIGI-HMM (Waack et al., 2006), INDeGenIUS (Shrivastava et al., 2010) and PAI-IDA (Tu and Ding, 2003). The subprograms were executed in a local system and the results were processed using the EGID algorithm (Che et al., 2011) to get a consensus result. IslandViewer is an integration of three subprograms: IslandPick, SIGI-HMM and IslandPath. The integrated result was downloaded from the IslandViewer server (www.pathogenomics.sfu.ca/islandviewer). Results of GIST and IslandViewer were further processed by an in-house Perl script to extract the genes included in the genomic islands. The putative HGT-derived genes predicted by Merhej et al. (2011) and by Le et al. (2012) were extracted from the original publications. Specifically, Merhej et al. (2011) identified 152 HGT-derived genes in the *R. felis* genome that are linked with organisms other than SFG *Rickettsia*. Le et al. (2012) identified 11 instances of HGT from outside Rickettsiales into the *R. felis* genome.

The predicted HGT-derived genes or genomic islands by different methods were spatially mapped to the *R. felis* genome and visualized in Geneious 6.0 (Biomatters). An “overlap factor” (OF) was employed as a criterion to compare the outcomes of different methods by assessing the overlap. This was expressed as the negative logarithm of the likelihood that the overlap was obtained by chance. To compute an OF, the number of the same genes predicted by each method pair was counted, and the OF was calculated following the probability mass function of the hypergeometric distribution:

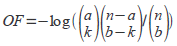

Where *n* is the total number of genes; *a* and *b* are the numbers of genes predicted by two methods, respectively; *k* is the number of genes overlapping by two sets of results. The larger an OF is, the more overlapping, and thus more consistent the two sets of results are, and the more likely it is that the two methods are identifying the same group of genes.

## Results and discussion

### Performance on simulated genomic data

#### Testing precision and recall

In all experimental groups under the idealized tree topology, a clear bimodal distribution was observed (Figure 2C), which is expected when HGT is present in the data. Meanwhile, none of the negative control groups have an identifiable *zero* peak, which is equivalent to a vertical history for all genes (no HGT events). Both cutoff criteria achieve high precision and recall simultaneously. In particular, under the conservative criterion, 99.4% of the prediction results are true positives. Meanwhile, they cover over 91.3% of all actual HGT-derived genes. The more relaxed criterion still achieves a precision of 95.3% and a recall of 96.8%.

In tests under randomized tree topologies, a larger transitional zone is present between peaks of the expected bimodal distribution, which is also frequently observed in real datasets. Both, precision and recall are affected compared to the idealized scenario, but in different measures. The conservative cutoff still maintains reasonably high precision (81.6%) and recall (90.5%) simultaneously, relative to the known number of HGT events. The relaxed criterion keeps equally high recall (96.6%) but its precision drops significantly (42.6%) (Figures 3 and S2).

**Figure 3.**
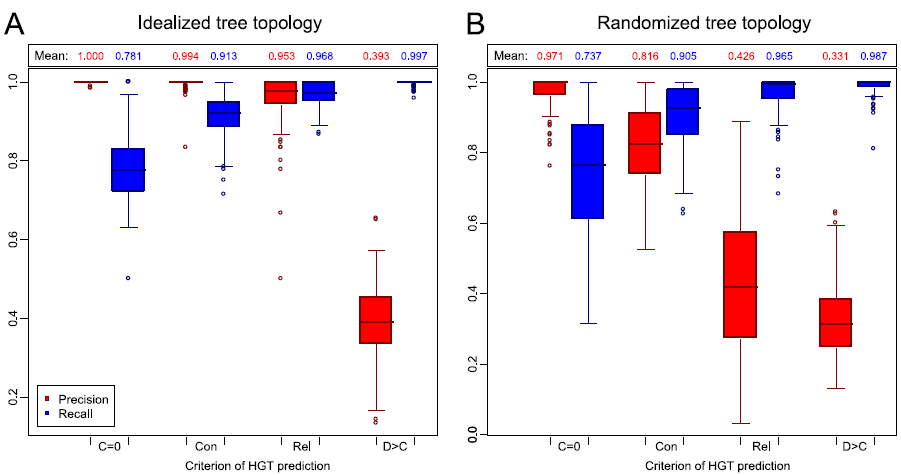
Comparison of performance of HGTector and conventional BLAST-based method on simulated genomes. The methods were tested on the simulated genomic data under an idealized **(A)** or randomized **(B)** tree topology. “Con” and “Rel” represent conservative and relaxed criteria of choosing cutoff in HGTector analysis. “C=0” and “D>C” are two criteria in the conventional BLAST-based method. Each experimental group is composed of 100 tests. Distribution of results in terms of precision (red) and recall (blue) is depicted by box plots. The mean value of each group is label above the corresponding column.

Because the test under randomized topologies is a better representation of real datasets, its result serve as a better reference for the practical consideration of HGTector application. Given the simultaneously high precision and recall, we recommend using the conservative cutoff for both initial HGT candidate screening (to be followed by phylogenetic analysis or other analyses) and standalone HGT discovery (when further in-depth analyses are not applicable). The relaxed cutoff may be considered only when the user wants to maximize discovery rate in an initial screening, in spite of its higher false positive rate. The comparison between idealized and randomized topologies further indicates the positive correlation between prediction success and a properly defined grouping scenario, in which: (1) the *self-close* clade is relatively distant from any other organisms; (2) there are multiple subclades in the *close* group, each having similar number of taxa represented in the database.

Varying global HGT rates seemingly show little effect on the stability of precision and recall of this method (Figure S3), suggesting that the predicting power is independent of HGT rate.

In comparison, the performance of both conventional BLAST-based approaches (C=0 and D>C) is notably unbalanced. Atypical genes falling under the C=0 criterion have the highest precision (100.0% and 97.1%, for idealized and randomized tree topology, respectively), but very low recall (78.1% and 73.7%). The D>C criterion has high recall (99.7% and 98.7%) but extremely low precision (39.3% and 33.1%), showing an intolerably high false positive rate (Figures 3 and S2). From a practical perspective, C=0 is too stringent, thus omitting a big portion of true HGT-derived genes affected by stochastic events, while D>C is too relaxed and not capable of differentiating genes that have high-score *distal* hits merely due to stochastic reasons instead of HGT (see below). It is to be mentioned that C=0 is not applicable to real datasets due to frequent genome annotation errors and ORFans, both of which may have a zero *close* weight.

#### Evaluating other evolutionary scenarios

The impact of other evolutionary events on the fingerprint and the division between typical and atypical gene populations was explored. Outgoing HGT (from *self* to *distal*) seemingly does not significantly alter the *close* weight of a gene. Gene loss decreases *close* weight and gene duplication increases it, both within an insignificant range (Figure 2C, D). Most importantly, genes within these three categories of evolutionary history still fall within the typical region and were not mistakenly detected as atypical by HGTector.

It is particularly notable that the majority of genes derived from ancient HGT events are located in the transitional zone (Figure 2D). Expectedly, they constitute a considerable portion of false positives in our analyses. In other words, depending on cutoff, more of them are likely to be placed in the typical population, instead of the atypical, and presumably non-vertical population. A similar pattern was observed for secondary HGT events. Although not frequent in the simulated genomes here, these events are actually very frequent in real datasets, as HGT frequency is higher between closely related organisms (Popa and Dagan, 2011). However, in the conventional BLAST method (D>C) most false positives are composed of mainly outgoing HGTs but only a few ancient HGTs.

### Application to real datasets

The *close* weight distributions for real datasets exhibit a bimodal distribution containing a broad *typical* region and a sharp atypical peak that is located close to zero (as expected) (Figures 4A-C and S4). Therefore, the cutoffs that divide genes with atypical BLAST hit distribution from typical ones can be set accordingly and HGT events can be predicted based on the cutoffs. As an example, fingerprint plots for *Rickettsia* and *Galdieria* exhibit an apparent overlapping pattern between the atypical peak and the HGT-derived genes identified by previous phylogenetic studies (Merhej et al., 2011; Schonknecht et al., 2013)(Figures S4E and S5D). In contrast, the human genome does not have an apparent atypical peak, which is also expected (Figure S4F).

**Figure 4.**
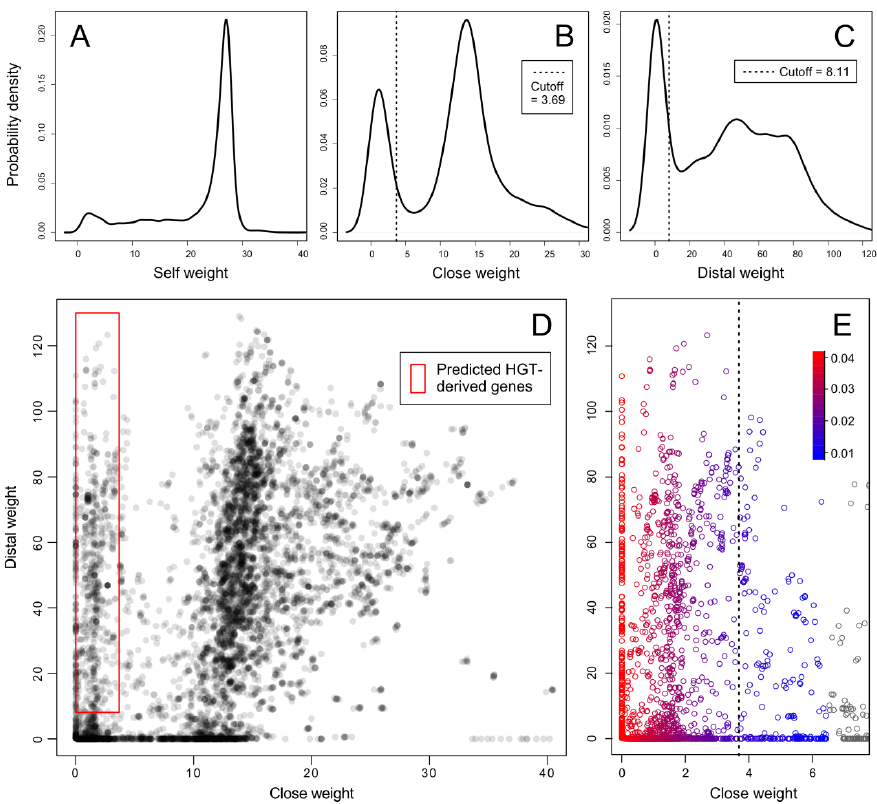
Fingerprint of seven *Rickettsia* genomes. BLAST hit weights of all protein-coding genes in the seven *Rickettsia* genomes are plotted. (**A-C**) Kernel density functions of the *self, close* and *distal* weights. The x-axis represents the weight of each gene. The y-axis represents the probability density of genes with the corresponding weight in the genomes. In this example, rule 1 (*close* weight < cutoff) and rule 2 (*distal* weight >= cutoff) were applied. The *close* and *distal* cutoffs computed under the conservative criterion are indicated by dashed lines. The values of the cutoffs are denoted in each panel. (**D**) A scatter plot of the *distal* weight against the *close* weight, showing the clustering pattern of the genes. Each dot represents one gene. Genes predicted to be HGT-derived are framed by a red rectangle. (**E**) A zoom-in view of the left part of the previous plot. Genes that fall within the atypical region in the *close* weight distribution are colored by a blue-red color scheme based on the density-based silhouette (dbs), a measure of confidence that this gene belongs to the atypical cluster of genes (red = high confidence). The *close* cutoff used in the subsequent analyses is indicated by a dashed line.

#### Analysis of the Rickettsia genomes

The global fingerprint describing the pattern of BLAST hit distribution of the *Rickettsia* analysis is illustrated in Figure 4A-C. Results from Hartigans’ dip test strongly support the non-unimodality of all three weight distributions (p-values < 2.2×10^−16^). The seven *Rickettsia* genomes contain a total number of 8484 annotated chromosomal protein-coding genes, of which 800 genes have an atypical *close* weight and a typical *distal* weight, and thus potentially HGT-derived (Tables 2 and S1). The percentage of putative HGT-derived genes in a genome ranges from 6.05% (76 genes) in *R. akari* to 18.29% (256 genes) in *R. felis*. The number of putative HGT-derived genes per genome is positively correlated to the size of the genome (R^2^ = 0.755), implying contribution of HGT to the relative genome expansions in the overall reductive trend of *Rickettsia* genome evolution, confirming previous studies (Merhej and Raoult, 2011). It is notable that the *R. felis* genome contains significantly more putative HGT-derived genes (256) than the rest of the genomes (90.7 ± 20.5, mean and standard deviation), suggesting a particular prevalence of potential HGT events in *R. felis* evolution. The comparison between the fingerprint on *R. felis* genome alone (Figure S5A-C) and the global fingerprint (Figure 4A-C) clearly reveals that *R. felis* has a much larger atypical peak in the *close* weight distribution. This reinforces outcomes from Merhej et al. (2011), which found that the frequency of cross-species bacterial recombination in *R. felis* had been underestimated.

**Table 2.** Summary of genes predicted to be horizontally acquired in seven *Rickettsia* genomes

| Species | Size of chromosome (Mb) | No. of genes | No. of predicted HGTs | % of predicted HGTs |
| --- | --- | --- | --- | --- |
| *R. akari* | 1.23 | 1256 | 76 | 6.05% |
| *R. felis* | 1.49 | 1400 | 256 | 18.29% |
| *R. massiliae* | 1.36 | 968 | 93 | 9.61% |
| *R. slovaca* | 1.28 | 1114 | 72 | 6.46% |
| *R. rickettsii* | 1.26 | 1342 | 98 | 7.30% |
| *R. africae* | 1.28 | 1030 | 78 | 7.57% |
| *R. conorii* | 1.27 | 1374 | 127 | 9.24% |
| **Mean** | 1.31 | 1212 | 114.3 | 9.43% |

To further explore the biological information behind the predicted patterns, the results are further summarized in three separate contexts: by putative donor group, by functional annotation and by gene orthology. Similar to previous in depth analyses by Merhej et al. (2011), our analyses revealed frequent donor links, such as Legionellales, Enterobacteriales and Burkholderiales, and frequently transferred gene categories, such as genes encoding transposases and genes involved in phage and plasmid activity (data not shown).

#### Stochastic manipulation of Rickettsia data: Stability of prediction

The results of these simulations (Figures S6 and S7) suggest that HGTector’s performance is notably insensitive to database error. Even under an extremely high proportion (10%) of mislabeled genes, its precision remains close to 100%. As a comparison, the precision of the conventional BLAST-based approach D>C drops below 15% (Figure S6A). Gene loss and rate variation in the *close* group also have remarkably weaker deleterious effects on precision of HGTector than that of conventional BLAST based methods under the D>C scenario (Figure S6B, C). Rate variations in input genomes destabilize prediction results at a certain range (5-15%), but overall, precision remains high in HGTector when rate increases, as compared to conventional BLAST (Figure S6D). Taxon sampling bias has little effect when it happens to distal organisms. However, in some close organisms (such as *R. prowazekii* and *R. bellii*), this effect may severely compromise the predicting power if the bias is strong (for example, a certain taxon group has many more representatives than others) (Figure S7).

The above results suggest that our method is generally unaffected by stochastic events, some of which are major challenges to current methods (see Introduction). The only concern is the taxon sampling bias in the *close* group, an issue that can be alleviated by properly defining grouping scenario and masking redundant taxonomic groups using HGTector’s flexible functionality (see Methods - Computational pipeline).

#### Comparison of HGTector with other methods for analysis of the R. felis genome

Together, HGTector and six other methods identified 595 putative HGT-derived genes, out of all 1400 chromosomal protein-coding genes in the *R. felis* genome (42.5%) (Table 2 and S2, Figure 5). Of these 595 genes, 82 have been uniquely identified by HGTector. There’s a considerable portion of overlapping hits between each pair of results, but none of the programs produce completely identical HGT predictions. As expected, there is relatively more overlap between two methods of the same category. Meanwhile, the overlap factors (OFs) between two methods from different categories are significantly lower. However, HGTector is a notable exception because its overlap with any other method of the three categories is significantly higher than between other methods from different categories (Table S3 and Figure S8).

**Figure 5.**
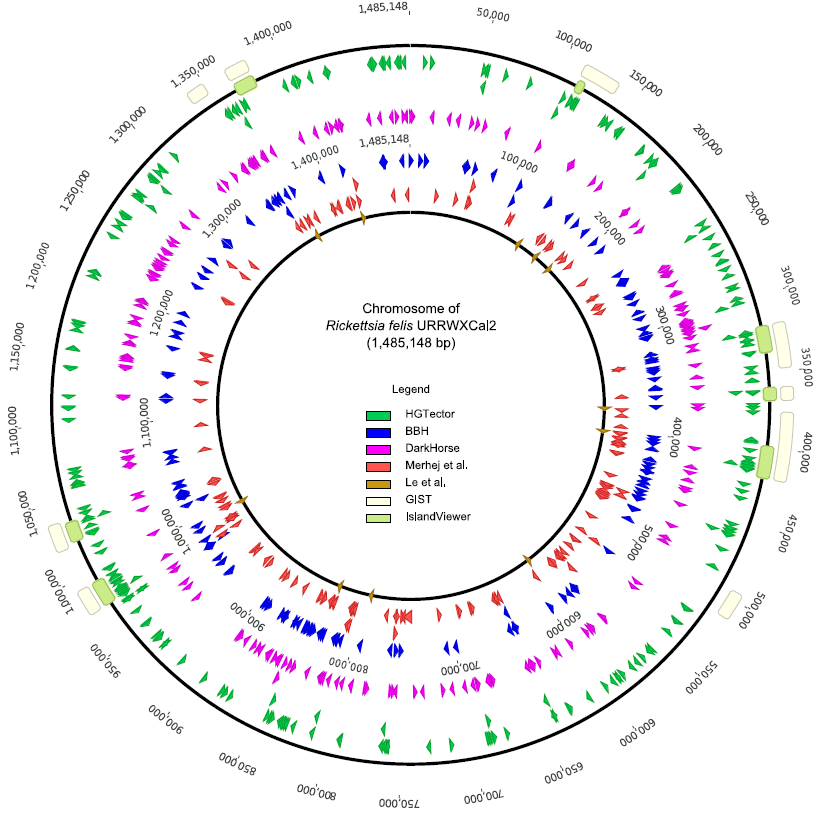
Predicted HGTs by multiple methods mapped onto the *Rickettsia felis* genome. A circular view of the whole chromosome of *R. felis*, with genomic islands (GIs) predicted by IslandViewer and GIST indicated by boxes, and putative HGT-derived genes predicted by other methods indicated by arrowheads.

By comparing the results of HGTector and the results of the other two BLAST-based methods, it is noticeable that HGTector can effectively exclude false positives caused by database errors. For instance, BBH and DarkHorse respectively predicted 61 and 97 genes to be acquired from *Ixodes scapularis* (deer tick), which is a common host of *Rickettsia*. Therefore, it is likely that the sequenced *Ixodes* samples contained *Rickettsia* endosymbionts, and its DNA was also sequenced and indiscriminately labeled as *Ixodes* DNA in the unassembled shotgun sequences (PRJNA34667). In other words, these are potential database errors. In contrast, in HGTector’s result, none of these genes were predicted as HGT-derived, because they have considerable amount of *close* hits (violation of rule 1), despite the presence of a single high-score *Ixodes* hit.

Detection of HGT is fraught with challenges, such as ambiguity in compositional features and phyletic patterns (Langille et al., 2008), difficulty in phylogenetic reconstruction (Philippe et al., 2011), interference from database error and incompleteness (Kohane et al., 2012). The observed low consistency between methods is not surprising, and has precedents in multiple previous studies (Lawrence and Ochman, 2002; Ragan et al., 2006; Zhaxybayeva, 2009). Given the significant overlap of our method with all other previously employed approaches, we suggest that our method is effective in producing meaningful prediction results on real datasets.

## Conclusion

In this paper a novel method for genome-wide detection of vertical versus non-vertical gene history (in particular, putative HGT events) is presented. It features a statistical analysis of BLAST hit distribution patterns in the context of a priori defined phylogenetic hierarchies. The innovation of this method is the systematic consideration of all BLAST hits of all genes within selected genomes. This is in contrast to conventional BLAST-based approaches, which typically rely on a single best hit for each gene (see Introduction). The three-category grouping scenario is a simplified but effective implementation of prior phylogenetic knowledge into a BLAST hit distribution analysis. The set-up allows high flexibility in group assignments that best match the taxonomic level of the user’s interest, as well as immediate response to frequent changes in microbial taxonomy. The most remarkable advantage of this systematic approach is that it captures the overall image of gene evolution while being immune to stochastic events. As demonstrated in this study, stochastic events such as gene loss, rate variation and database error may impose serious problem to conventional methods (see Introduction), but have comparatively negligible effects on HGTector.

The core assumption of the method is that the typical and atypical parts of the BLAST hit distribution are distinguishable. This assumption is repeatedly supported by tests on both simulated and real-world genomic data (Figures 2, 4 and S4). With the proposed procedures of computing cutoffs and the rules of targeting genes that are likely to be HGT-derived, remarkable prediction success was achieved (Figures 3 and 5, Table S3). Given these results, we suggest that HGTector is a useful addition to conventional BLAST-based approaches.

The HGTector pipeline has advantages of speed, automation, compatibility and low requirement for computational resources, making this program a generally applicable tool for discovery of vertical, and non-vertical history of genes, as well as initial HGT prediction. It is especially suitable for gaining a rapid and comprehensive overview of newly sequenced genomes to identify their evolutionary and ecological linkages with other organisms, facilitating further exploration of the functional drivers of the dynamics of genome evolution. It has to be made clear that an atypical BLAST hit distribution is an empirical observation, rather than a strict certification of HGT. Since an HGT predominantly reflects a past evolutionary event, it is theoretically impossible to identify exact gene donors and mechanisms, and any analysis is only an approximation to possible scenarios. Therefore, we recommend HGTector as a discovery tool for a detection of potential HGT-derived genes that can be further analyzed with phylogenetic approaches. This is much more effective approach than a very time-consuming and technically challenging process of a priori phylogenetic analysis of all genes within all target genomes, which becomes decreasingly feasible as more genomic data are present.

## Acknowledgements

This research was funded by NSF DEB 1213740, awarded to KD.

## Supplementary material

File S1 Supplementary figures S1-S8.

File S2 Supplementary tables S1-S3.

## Authors’ contributions

QZ and KD conceived and designed the study. QZ developed the algorithm, wrote the program, performed the data analyses and drafted the manuscript. KD contributed to the design of algorithm and program, supervised the data analyses and edited the manuscript. MK contributed to the discussion and helped to revise the manuscript. All authors read and approved the final manuscript.

## References

Abby, S.S., Tannier, E., Gouy, M., and Daubin, V. (2012). Lateral gene transfer as a support for the tree of life. Proc Natl Acad Sci U S A 109, 4962–4967.

Altschul, S.F., Madden, T.L., Schäffer, A.A., Zhang, J., Zhang, Z., Miller, W., and Lipman, D.J. (1997). Gapped BLAST and PSI-BLAST: a new generation of protein database search programs. Nucleic Acids Res 25, 3389–3402.

Andam, C.P., Williams, D., and Gogarten, J.P. (2010). Natural taxonomy in light of horizontal gene transfer. Biol Philos 25.

Bapteste, E., O’Malley, M.A., Beiko, R.G., Ereshefsky, M., Gogarten, J.P., Franklin-Hall, L., Lapointe, F.-J., Dupré, J., Dagan, T., Boucher, Y., et al. (2009). Prokaryotic evolution and the tree of life are two different things. Biol Direct 4, 34.

Biomatters Geneious version 6.0. Available from http://www.geneious.com/.

Blanc, G., Ogata, H., Robert, C., Audic, S., Claverie, J.-M., and Raoult, D. (2007). Lateral gene transfer between obligate intracellular bacteria: evidence from the Rickettsia massiliae genome. Genome Res 17, 1657–1664.

Boto, L. (2010). Horizontal gene transfer in evolution: facts and challenges. Proc Biol Sci 277, 819–827.

Brenner, S.E. (1999). Errors in genome annotation. Trends Genet 15, 132–133.

Castresana, J. (2000). Selection of conserved blocks from multiple alignments for their use in phylogenetic analysis. Mol Biol Evol 17, 540–552.

Charlebois, R.L., Clarke, G.D., Beiko, R.G., and St Jean, A. (2003). Characterization of species-specific genes using a flexible, web-based querying system. FEMS Microbiol Lett 225, 213–220.

Charlebois, R.L., and Doolittle, W.F. (2004). Computing prokaryotic gene ubiquity: rescuing the core from extinction. Genome Res 14, 2469–2477.

Che, D., Hasan, M.S., Wang, H., Fazekas, J., Huang, J., and Liu, Q. (2011). EGID: an ensemble algorithm for improved genomic island detection in genomic sequences. Bioinformation 7, 311–314.

Chen, F., Mackey, A.J., Stoeckert, C.J., Jr., and Roos, D.S. (2006). OrthoMCL-DB: querying a comprehensive multi-species collection of ortholog groups. Nucleic Acids Res 34, D363–368.

Clarke, G.D.P., Beiko, R.G., Ragan, M.A., and Charlebois, R.L. (2002). Inferring genome trees by using a filter to eliminate phylogenetically discordant sequences and a distance matrix based on mean normalized BLASTP scores. J Bacteriol 184, 2072–2080.

Conesa, A., Götz, S., García-Gómez, J.M., Terol, J., Talón, M., and Robles, M. (2005). Blast2GO: a universal tool for annotation, visualization and analysis in functional genomics research. Bioinformatics 21, 3674–3676.

Dagan, T., Artzy-Randrup, Y., and Martin, W. (2008). Modular networks and cumulative impact of lateral transfer in prokaryote genome evolution. Proc Natl Acad Sci U S A 105, 10039–10044.

Darriba, D., Taboada, G.L., Doallo, R., and Posada, D. (2012). jModelTest 2: more models, new heuristics and parallel computing. Nat Methods 9, 772–772.

Delsuc, F., Brinkmann, H., and Philippe, H. (2005). Phylogenomics and the reconstruction of the tree of life. Nat Rev Genet 6, 361–375.

Doolittle, W.F. (1999). Phylogenetic classification and the universal tree. Science 284, 2124–2129.

Doolittle, W.F., Boucher, Y., Nesbø, C.L., Douady, C.J., Andersson, J.O., and Roger, A.J. (2003). How big is the iceberg of which organellar genes in nuclear genomes are but the tip? Philos Trans R Soc Lond B Biol Sci 358, 39–57; discussion 57–58.

Eisen, J.A. (2000). Assessing evolutionary relationships among microbes from whole-genome analysis. Curr Opin Microbiol 3, 475–480.

Fischer, D., and Eisenberg, D. (1999). Finding families for genomic ORFans. Bioinformatics 15, 759–762.

Frederic, I., Philippe, G., and Michele, E. (2013). Pastecs: package for analysis of space-time ecological series. R package version 1.3-13.

Frickey, T., and Lupas, A.N. (2004). PhyloGenie: automated phylome generation and analysis. Nucleic Acids Res 32, 5231–5238.

Gouy, M., Guindon, S., and Gascuel, O. (2010). SeaView version 4: A multiplatform graphical user interface for sequence alignment and phylogenetic tree building. Mol Biol Evol 27, 221–224.

Hacker, J., and Kaper, J.B. (2000). Pathogenicity islands and the evolution of microbes. Annu Rev Microbiol 54, 641–679.

Hartigan, J.A., and Hartigan, P. (1985). The dip test of unimodality. The Annals of Statistics, 70–84.

Hasan, M.S., Liu, Q., Wang, H., Fazekas, J., Chen, B., and Che, D. (2012). GIST: Genomic island suite of tools for predicting genomic islands in genomic sequences. Bioinformation 8, 203–205.

Hsiao, W., Wan, I., Jones, S.J., and Brinkman, F.S.L. (2003). IslandPath: aiding detection of genomic islands in prokaryotes. Bioinformatics 19, 418–420.

Kendall, M.G. (1976). Time-series (London: Griffin).

Kohane, I.S., Hsing, M., and Kong, S.W. (2012). Taxonomizing, sizing, and overcoming the incidentalome. Genet Med 14, 399–404.

Koonin, E.V., Makarova, K.S., and Aravind, L. (2001). Horizontal gene transfer in prokaryotes: quantification and classification. Annu Rev Microbiol 55, 709–742.

Koonin, E.V., and Wolf, Y.I. (2009). The fundamental units, processes and patterns of evolution, and the tree of life conundrum. Biol Direct 4, 33.

Koski, L.B., and Golding, G.B. (2001). The closest BLAST hit is often not the nearest neighbor. J Mol Evol 52, 540–542.

Langille, M.G.I., and Brinkman, F.S.L. (2009). IslandViewer: an integrated interface for computational identification and visualization of genomic islands. Bioinformatics 25, 664–665.

Langille, M.G.I., Hsiao, W.W.L., and Brinkman, F.S.L. (2008). Evaluation of genomic island predictors using a comparative genomics approach. BMC Bioinformatics 9, 329.

Lawrence, J.G., and Ochman, H. (1997). Amelioration of bacterial genomes: rates of change and exchange. J Mol Evol 44, 383–397.

Lawrence, J.G., and Ochman, H. (2002). Reconciling the many faces of lateral gene transfer. Trends Microbiol 10, 1–4.

Le, P.T., Golaconda Ramulu, H., Guijarro, L., Paganini, J., Gouret, P., Chabrol, O., Raoult, D., and Pontarotti, P. (2012). An automated approach for the identification of horizontal gene transfers from complete genomes reveals the rhizome of Rickettsiales. BMC Evol Biol 12, 243.

Long, M., Betrán, E., Thornton, K., and Wang, W. (2003). The origin of new genes: glimpses from the young and old. Nat Rev Genet 4, 865–875.

Maddison, D.R., Swofford, D.L., and Maddison, W.P. (1997). NEXUS: an extensible file format for systematic information. Syst Biol 46, 590–621.

Maechler, M., and Ringach, D. (2009). Diptest: Hartigan’s dip test statistic for unimodality-corrected code. R package version 025-2.

McDaniel, L.D., Young, E., Delaney, J., Ruhnau, F., Ritchie, K.B., and Paul, J.H. (2010). High frequency of horizontal gene transfer in the oceans. Science 330, 50.

Menardi, G. (2010). Density-based Silhouette diagnostics for clustering methods. Statistics and Computing 21, 295–308.

Merhej, V., Notredame, C., Royer-Carenzi, M., Pontarotti, P., and Raoult, D. (2011). The rhizome of life: the sympatric Rickettsia felis paradigm demonstrates the random transfer of DNA sequences. Mol Biol Evol 28, 3213–3223.

Merhej, V., and Raoult, D. (2011). Rickettsial evolution in the light of comparative genomics. Biol Rev Camb Philos Soc 86, 379–405.

Moreno-Hagelsieb, G., and Latimer, K. (2008). Choosing BLAST options for better detection of orthologs as reciprocal best hits. Bioinformatics 24, 319–324.

Nelson, K.E., Clayton, R.A., Gill, S.R., Gwinn, M.L., Dodson, R.J., Haft, D.H., Hickey, E.K., Peterson, J.D., Nelson, W.C., Ketchum, K.A., et al. (1999). Evidence for lateral gene transfer between Archaea and bacteria from genome sequence of Thermotoga maritima. Nature 399, 323–329.

Ochman, H., Lawrence, J.G., and Groisman, E.A. (2000). Lateral gene transfer and the nature of bacterial innovation. Nature 405, 299–304.

Ogata, H., Renesto, P., Audic, S., Robert, C., Blanc, G., Fournier, P.-E., Parinello, H., Claverie, J.-M., and Raoult, D. (2005). The genome sequence of Rickettsia felis identifies the first putative conjugative plasmid in an obligate intracellular parasite. PLoS Biol 3, e248.

Overbeek, R., Fonstein, M., D’Souza, M., Pusch, G.D., and Maltsev, N. (1999). The use of gene clusters to infer functional coupling. Proc Natl Acad Sci U S A 96, 2896–2901.

Philippe, H., Brinkmann, H., Lavrov, D.V., Littlewood, D.T.J., Manuel, M., Wörheide, G., and Baurain, D. (2011). Resolving difficult phylogenetic questions: why more sequences are not enough. PLoS Biol 9, e1000602.

Podell, S., and Gaasterland, T. (2007). DarkHorse: a method for genome-wide prediction of horizontal gene transfer. Genome Biol 8, R16.

Podell, S., Gaasterland, T., and Allen, E.E. (2008). A database of phylogenetically atypical genes in archaeal and bacterial genomes, identified using the DarkHorse algorithm. BMC Bioinformatics 9, 419.

Popa, O., and Dagan, T. (2011). Trends and barriers to lateral gene transfer in prokaryotes. Curr Opin Microbiol 14, 615–623.

R Core Team (2013). R: A Language and Environment for Statistical Computing (Vienna, Austria).

Ragan, M.A., Harlow, T.J., and Beiko, R.G. (2006). Do different surrogate methods detect lateral genetic transfer events of different relative ages? Trends Microbiol 14, 4–8.

Rambaut, A. (2013). FigTree.

Schonknecht, G., Chen, W.H., Ternes, C.M., Barbier, G.G., Shrestha, R.P., Stanke, M., Brautigam, A., Baker, B.J., Banfield, J.F., Garavito, R.M., et al. (2013). Gene transfer from bacteria and archaea facilitated evolution of an extremophilic eukaryote. Science 339, 1207–1210.

Shokralla, S., Spall, J.L., Gibson, J.F., and Hajibabaei, M. (2012). Next-generation sequencing technologies for environmental DNA research. Mol Ecol 21, 1794–1805.

Shrivastava, S., Reddy, C.V.S.K., and Mande, S.S. (2010). INDeGenIUS, a new method for high-throughput identification of specialized functional islands in completely sequenced organisms. J Biosci (Bangalore) 35, 351–364.

Sicheritz-Pontén, T., and Andersson, S.G. (2001). A phylogenomic approach to microbial evolution. Nucleic Acids Res 29, 545–552.

Silverman, B.W. (1981). Using Kernel density estimates to investigate multimodality. J R Statist Soc B 43, 97–99.

Silverman, B.W. (1986). Density Estimation (London: Chapman and Hall).

Simonson, A.B., Servin, J.A., Skophammer, R.G., Herbold, C.W., Rivera, M.C., and Lake, J.A. (2005). Decoding the genomic tree of life. Proc Natl Acad Sci U S A 102 Suppl 1, 6608–6613.

Smillie, C.S., Smith, M.B., Friedman, J., Cordero, O.X., David, L.A., and Alm, E.J. (2011). Ecology drives a global network of gene exchange connecting the human microbiome. Nature 480, 241–244.

Stamatakis, A. (2006). RAxML-VI-HPC: maximum likelihood-based phylogenetic analyses with thousands of taxa and mixed models. Bioinformatics 22, 2688–2690.

Stanhope, M.J., Lupas, A., Italia, M.J., Koretke, K.K., Volker, C., and Brown, J.R. (2001). Phylogenetic analyses do not support horizontal gene transfers from bacteria to vertebrates. Nature 411, 940–944.

Tatusov, R.L., Koonin, E.V., and Lipman, D.J. (1997). A genomic perspective on protein families. Science 278, 631–637.

Thompson, J.D., Higgins, D.G., and Gibson, T.J. (1994). CLUSTAL W: improving the sensitivity of progressive multiple sequence alignment through sequence weighting, position-specific gap penalties and weight matrix choice. Nucleic Acids Res 22, 4673–4680.

Tu, Q., and Ding, D. (2003). Detecting pathogenicity islands and anomalous gene clusters by iterative discriminant analysis. FEMS Microbiol Lett 221, 269–275.

Vernikos, G.S., and Parkhill, J. (2006). Interpolated variable order motifs for identification of horizontally acquired DNA: revisiting the Salmonella pathogenicity islands. Bioinformatics 22, 2196–2203.

Vitorino, L., Chelo, I.M., Bacellar, F., and Zé-Zé, L. (2007). Rickettsiae phylogeny: a multigenic approach. Microbiology 153, 160–168.

Waack, S., Keller, O., Asper, R., Brodag, T., Damm, C., Fricke, W.F., Surovcik, K., Meinicke, P., and Merkl, R. (2006). Score-based prediction of genomic islands in prokaryotic genomes using hidden Markov models. BMC Bioinformatics 7, 142.

Weinert, L.A., Welch, J.J., and Jiggins, F.M. (2009). Conjugation genes are common throughout the genus Rickettsia and are transmitted horizontally. Proc Biol Sci 276, 3619–3627.

Willerslev, E., Mourier, T., Hansen, A.J., Christensen, B., Barnes, I., and Salzberg, S.L. (2002). Contamination in the draft of the human genome masquerades as lateral gene transfer. DNA Sequence 13, 75–76.

Wolf, Y.I., Aravind, L., and Koonin, E.V. (1999). Rickettsiae and Chlamydiae: evidence of horizontal gene transfer and gene exchange. Trends Genet 15, 173–175.

Wong, W.C., Maurer-Stroh, S., and Eisenhaber, F. (2010). More than 1,001 problems with protein domain databases: transmembrane regions, signal peptides and the issue of sequence homology. PLoS Comput Biol 6, e1000867.

Zhaxybayeva, O. (2009). Detection and quantitative assessment of horizontal gene transfer. Methods Mol Biol 532, 195–213.

Zhaxybayeva, O., and Doolittle, W.F. (2011). Lateral gene transfer. Curr Biol 21, R242–246.

Zhaxybayeva, O., Nesbø, C.L., and Doolittle, W.F. (2007). Systematic overestimation of gene gain through false diagnosis of gene absence. Genome Biol 8, 402.

